# FGF signalling regulates enhancer activation during ear progenitor induction

**DOI:** 10.1101/595058

**Authors:** Monica Tambalo, Maryam Anwar, Mohi Ahmed, Andrea Streit

**Affiliations:** Centre for Craniofacial and Regenerative Biology, Faculty of Dental, Oral and Craniofacial Sciences, King’s College London, London SE1 9RT, UK

**Keywords:** Cranial ganglia, cis-regulatory elements, ear, gene networks, hearing, placode

## Abstract

The fibroblast growth factor pathway is essential for inner ear induction in many vertebrates, however how it regulates the chromatin landscape to coordinate the activation of otic genes remains unclear. Here we show that FGF exposure of sensory progenitors leads to rapid deposition of active chromatin marks H3K27ac near hundreds of FGF-responsive, otic-epibranchial progenitor (OEP) genes, while H3K27ac is depleted in the vicinity of non-otic genes. Genomic regions that gain H3K27ac act as cis-regulatory elements controlling OEP gene expression in time and space and define a unique transcription factor signature likely to control their activity. Finally, we provide evidence that in response to FGF signalling the transcription factor dimer AP1 recruits the histone acetyl transferase p300 to OEP enhancers and that de novo acetylation is required for subsequent expression of OEP genes. Thus, during ear induction FGF signalling modifies the chromatin landscape to promote enhancer activation and chromatin accessibility.

## INTRODUCTION

The vertebrate inner ear is critical to relay auditory and vestibular information from the environment to the brain. Developmental malformations or postnatal damage of inner ear cells lead to permanent sensory defects as the ear does not have the ability to regenerate or repair. Despite extensive studies, our understanding of normal ear development has not yet translated into new biological approaches to alleviate such disorders or to promote sensory cell regeneration *in vivo*. This is partly due to the lack of mechanistic information downstream of signalling pathways that control ear development and of how signalling events are translated into changes in gene expression and cell behaviour.

The Fibroblast Growth Factor (FGF) pathway is crucially important for many steps in ear formation. During early development, FGF signalling mediates the induction of otic-epibranchial progenitors (OEPs) from a pool of sensory stem cells (Ladher et al., 2000; Maroon et al., 2002; Park and Saint-Jeannet, 2008; Phillips et al., 2001; Sun et al., 2007; Urness et al., 2010; Wright and Mansour, 2003; Yang et al., 2013a). Otic precursors segregate from epibranchial progenitors to form a transient structure, the otic placode, and subsequently the otic vesicle, which gives rise to the entire inner ear including the cochlear-vestibular ganglion (Chen and Streit, 2013; Groves and Fekete, 2012; Ladher, 2017; Ohyama et al., 2007; Whitfield, 2015). During these stages, FGF signalling promotes otic neurogenesis and mediates patterning events that subdivide the otic vesicle into distinct functional domains (Abello et al., 2010; Adamska et al., 2001; Alsina et al., 2004; Alvarez et al., 2003; Hammond and Whitfield, 2011; Leger and Brand, 2002; Maier and Whitfield, 2014). As the vesicle gradually acquires the shape of the adult ear, sensory patches emerge that generate auditory and vestibular sensory hair cells and their surrounding supporting cells. Within the cochlea, these cells form a stereotypical pattern that is critical for normal hearing, with FGF signalling controlling both their arrangement and their differentiation (Doetzlhofer et al., 2009; Haque et al., 2016; Mueller et al., 2002; Shim et al., 2005). Finally, FGFs have also been implicated in hair cell survival in adult ears as well as controlling their regeneration from supporting cells (Jiang et al., 2018; Kniss et al., 2016; Lee et al., 2016; Lush et al., 2019).

Upon ligand binding FGF receptors initiate different intracellular cascades including the ERK/MAPK pathway (Ornitz and Itoh, 2015) to regulate gene expression and subsequent changes in cell behaviour. Signalling information is integrated by non-coding regulatory enhancer regions to activate or repress downstream targets (Banerji et al., 1981; Catarino and Stark, 2018; Long et al., 2016; Shlyueva et al., 2014). Active enhancers are characterised by low nucleosome density (Boyle et al., 2008), recruitment of the histone acetylase p300 (Visel et al., 2009), acetylation of histone 3 lysine-27 (H3K27ac) in the enhancer-flanking nucleosomes (Creyghton et al., 2010; Kharchenko et al., 2011; Rada-Iglesias et al., 2012; Rada-Iglesias et al., 2011; Zentner et al., 2011), and the expression of RNA transcripts (Kim et al., 2010; Wang et al., 2011). Downstream of FGF signalling, different mechanisms regulate target gene expression. On one hand, Erk1/2 MAP-kinases can directly modify nucleosome forming histones and change epigenetic marks (Chen et al., 2007; Soloaga et al., 2003), and their inhibition can increase chromatin accessibility (Patel et al., 2013; Semprich et al., 2019). On the other hand, MAP kinases phosphorylate Ets transcription factors (e.g. Etv4 and Etv5) or the dimer forming leucine zippers cFos and cJun (AP1) (Gruda et al., 1994; Neuberg et al., 1989; Ornitz and Itoh, 2015; Tsang and Dawid, 2004) (for review: (Yang et al., 2003, 2013b), which then bind to enhancers and together with other factors, activate target genes. For example, AP1 cooperates with cell type specific transcription factors to recruit the BAF complex, which in turn remodels nucleosomes to establish a permissive chromatin state (Vierbuchen et al., 2017). Thus, modulation of FGF signalling appears to control changes in the epigenetic signature. However, in the context of *in vivo* ear development, the molecular mechanisms that translate FGF signalling into rapid transcriptional changes remain to be elucidated.

Here we study chromatin changes in response to FGF signalling during the earliest step of ear development, the induction of otic-epibranchial progenitors. We find that FGF stimulation of sensory precursors leads to deposition of H3K27ac marks in the vicinity of ear-specific, FGF-response genes and that these genomic regions act as ear-specific enhancers. *De novo* acetylation of H3K27 is required for otic gene expression and fate allocation. We show that AP1 plays a key role in this process: upon FGF signalling, AP1 recruits the histone acetylase p300 to ear enhancers, which in turn promotes H3K27 acetylation associated with increased chromatin accessibility and enhancer activation. Together these findings highlight that during ear induction, the initial response to Erk/MAPK signalling directly activates ear-specific enhancers, providing a molecular mechanism for rapid activation of gene expression downstream of FGF. In turn, these observations may impact on a variety of diseases and developmental disorders where FGFs play a major role.

## RESULTS

### FGF signalling induces dynamic changes in H3K27 acetylation

FGF signalling is critical to initiate the ear programme. Loss of FGFs or pathway inhibition results in the complete absence of ear precursors, while exposure of sensory progenitors to FGF induces otic epibranchial progenitors (OEPs) (Ladher et al., 2000; Maroon et al., 2002; Park and Saint-Jeannet, 2008; Phillips et al., 2001; Sun et al., 2007; Urness et al., 2010; Wright and Mansour, 2003; Yang et al., 2013a). Only few FGF targets have been identified, and the downstream epigenetic changes are unknown (Anwar et al., 2017; Yang et al., 2013a). To define the initial events as sensory progenitor cells are induced to become OEPs, we assessed changes in Histone 2 lysine 27 acetylation (H3K27ac) and trimethylation (H3K27me3) in response to FGF signalling. Histone acetylation is associated with increased accessibility of chromatin and gene activation, marking active enhancers and promoters, while H3K27me3 is associated with gene repression (for review see: Maston et al., 2012, Calo and Wysocka, 2013, Kimura, 2013). Sensory progenitors from HH6 chick embryos were cultured in the presence or absence of FGF2 for 6 hrs and processed for chromatin immunoprecipitation for H3K27ac and H3K27me3 followed by sequencing (nano-ChIP-seq; Figure 1A; (Adli and Bernstein, 2011). In both control and FGF-treated cells, transcription start sites (TSS) show H3K27ac accumulation and H3K27me3 depletion (Figure 1B). However, genome-wide comparison of both samples reveals significant changes in the distribution of H3K27ac marks.

**Figure 1.**
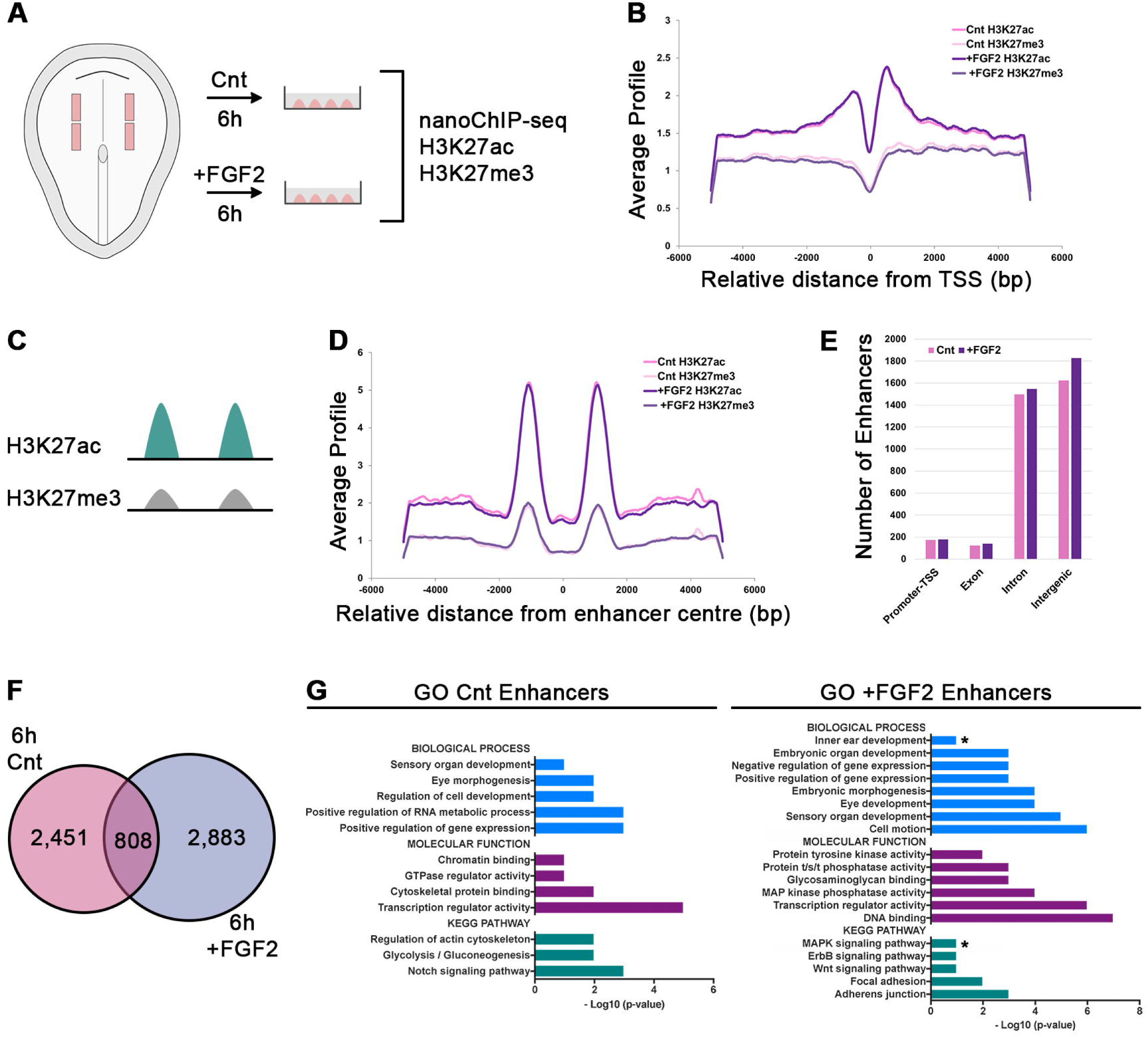
Distinct regulatory element usage during OEP induction by FGF2. (A) posterior sensory progenitor explants were dissected from 0ss (HH6) embyros and cultured in the presence (+FGF2) or absence (Cnt) of FGF2 for 6 hrs. Around 100 tissues per condition were processed for nanoChIP-seq. (B) Average read density of H3K27ac and H3K27me3 in control (pink) and +FGF2 treated cells (violet) around the transcription start site (TSS). Read densities are highest close to the TSS and display a peak-dip-peak pattern where the dip-region may represent bound transcriptional activators or repressors. (C) Bimodal distribution of H3K27ac (green) and absence/low H3K27me3 (grey) was used to identify enhancers. (D) In control (pink) and +FGF2 treated explants (violet), the average read density of H3K27ac and H3K27me3 around the centre of putative enhancers exhibit a bimodal distribution. H3K27me3 signals in both control and experimental cells is low, while H3K27ac is high as expected for active enhancers. (E) Bimodal H3K27ac peaks in control (Cnt; pink) and +FGF2 (violet) treated samples show a similar distribution with an average of around 170 located close to promotors, 130 in exonic, 1500 in intronic and 1700 in intergenic regions. (F) Venn-diagram representing putative enhancer elements unique in control (pink; 2,451) and +FGF2-treated sensory progenitors (violet; 2,883). 808 putative enhancers are shared in both conditions. (G) Putative enhancers were associated to the closest TSS. Gene ontology annotation reveals significant enrichment of ear and MAPK signalling related terms in enhancer associated genes in FGF2-treated sensory progenitors, but not in controls.

Association of H3K27ac peaks to the nearest TSS reveals that genes normally expressed in OEPs like the direct FGF targets *Spry1* and *Spry2*, the FGF-responsive transcript *Foxi3*, the transcription factor *Hesx1* and the chemokine *Cxcl14* show a marked gain in H3K27ac (Figure 2A, D; Supplementary Figure 1; Supplementary Figure 2A). In contrast, transcripts normally absent from OEPs and repressed by FGF signalling like *Pax6, Dlx5/6* and *Gata3* show a significant loss (Figure 2G; Supplementary Figure 6B-C), while there is no change in H3K27ac near sensory progenitor genes like *Eya2* (Figure 2H), whose expression precedes OEP induction by FGF (Ahrens and Schlosser, 2005; Litsiou et al., 2005; Pandur and Moody, 2000; Sato et al., 2010). Thus, changes in histone acetylation reflect changes in gene expression as sensory progenitors are specified as otic-epibranchial progenitors.

**Figure 2.**
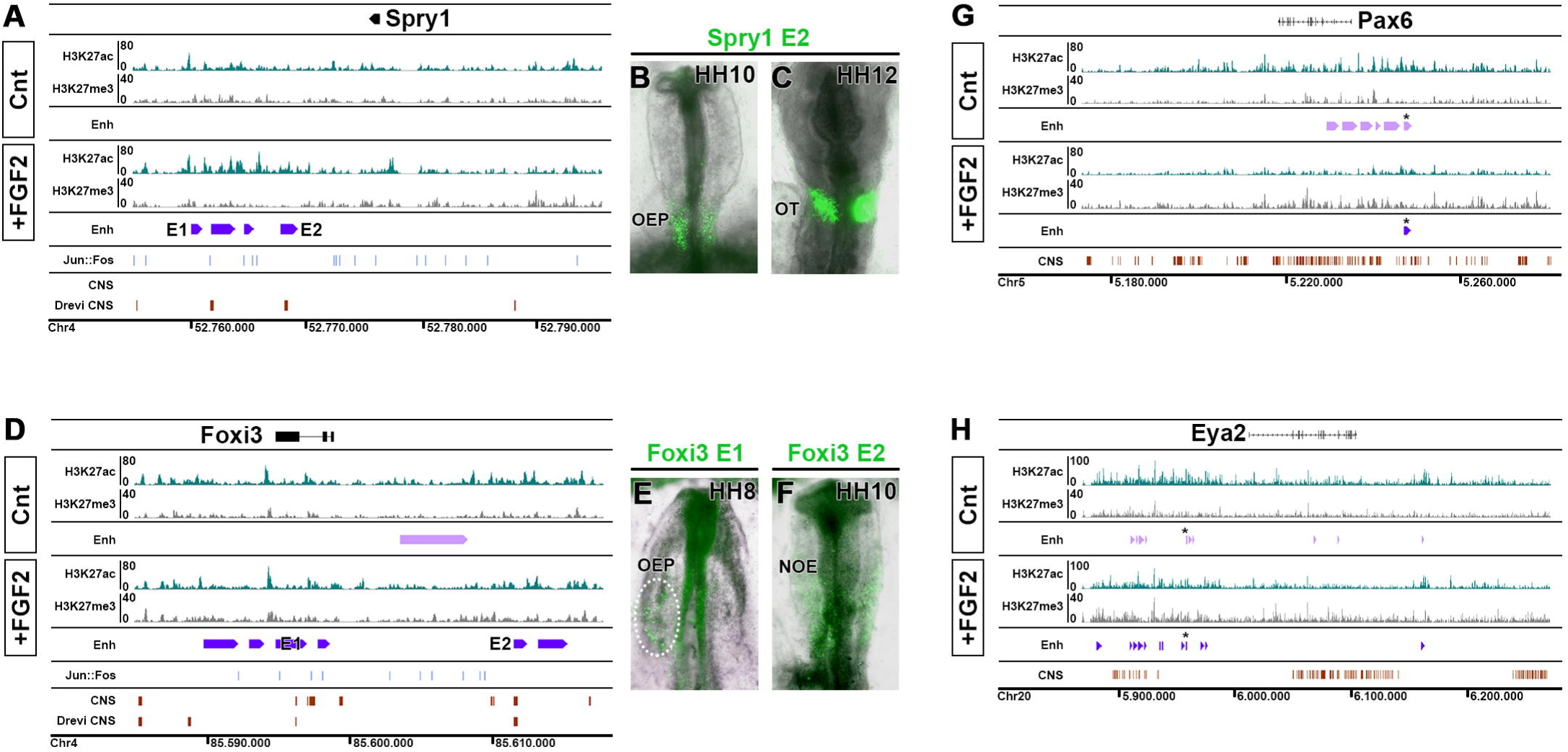
Ear specific enhancers gain H3K27ac upon FGF stimulation of sensory progenitors. IGB browser view of the Spry1 and Foxi3 locus (A, D), both expressed in OEPs, and of the FGF-repressed gene Pax6 (G) and the sensory progenitor gene Eya2 (H). H3K27ac tracks are shown in green and H3K27me3 tracks in grey. Enh: putative enhancers called by Homer and MACS2 (+FGF2 violet bars; control pink bars). Blue bars: Fos::Jun: location of AP1 binding motifs. CNS: conservation from PECAN alignments (ENSEMBL) and DREIVe are shown in red. (B, C) Spry1-E2 is active in OEPs and the otic cup. (E) Foxi3-E1 is active in OEPs at the 4-5ss (HH8), while Foxi3-E2 is activated later at the 10ss (HH10) in the non-otic ectoderm (NOE; F). (G) H3K27ac is lost in the Pax6 locus upon FGF treatment of sensory progenitor cells, while H3K27me3 is increased. The number of putative enhancers is reduced. (H) There are subtle changes in H3K27ac and H3K27me3 in the Eya2 locus. * marks an enhancer shared between controls and FGF2-treated cells.

### H3K27ac marks FGF-responsive enhancers in OEPs

Activation of gene expression is controlled by cis-regulatory elements and active enhancers are normally flanked by H3K27ac peaks (Figure 1C) (for review see: Maston et al., 2012, Calo and Wysocka, 2013, Kimura, 2013). Therefore, the increase in H3K27ac upon FGF stimulation may indicate the activation of ear enhancers. To identify putative enhancers genome-wide, we extracted bimodal H3K27ac peaks with a maximum distance of 3kb (Figure 1D) in sensory progenitors (control) and FGF-induced OEPs; these are mostly located in intronic or intergenic regions (Figure 1E). We find 2451 putative enhancers that are unique to control cells, 2883 specific to FGF2-treated progenitors, and 808 are common to both (Figure 1F, Supplementary Figure 3). FGF-induced putative enhancers preferentially associate with genes enriched in otic progenitors (from Chen et al., 2017): comparing all OEP transcripts to those with putative active enhancers shows significantly higher expression levels of the latter (Supplementary Figure 4). In addition, GO term analysis confirms that putative enhancer associated genes are linked to terms like inner ear development and MAP kinase signalling, while these terms are absent from genes in controls (Figure 1G).

**Figure 3.**
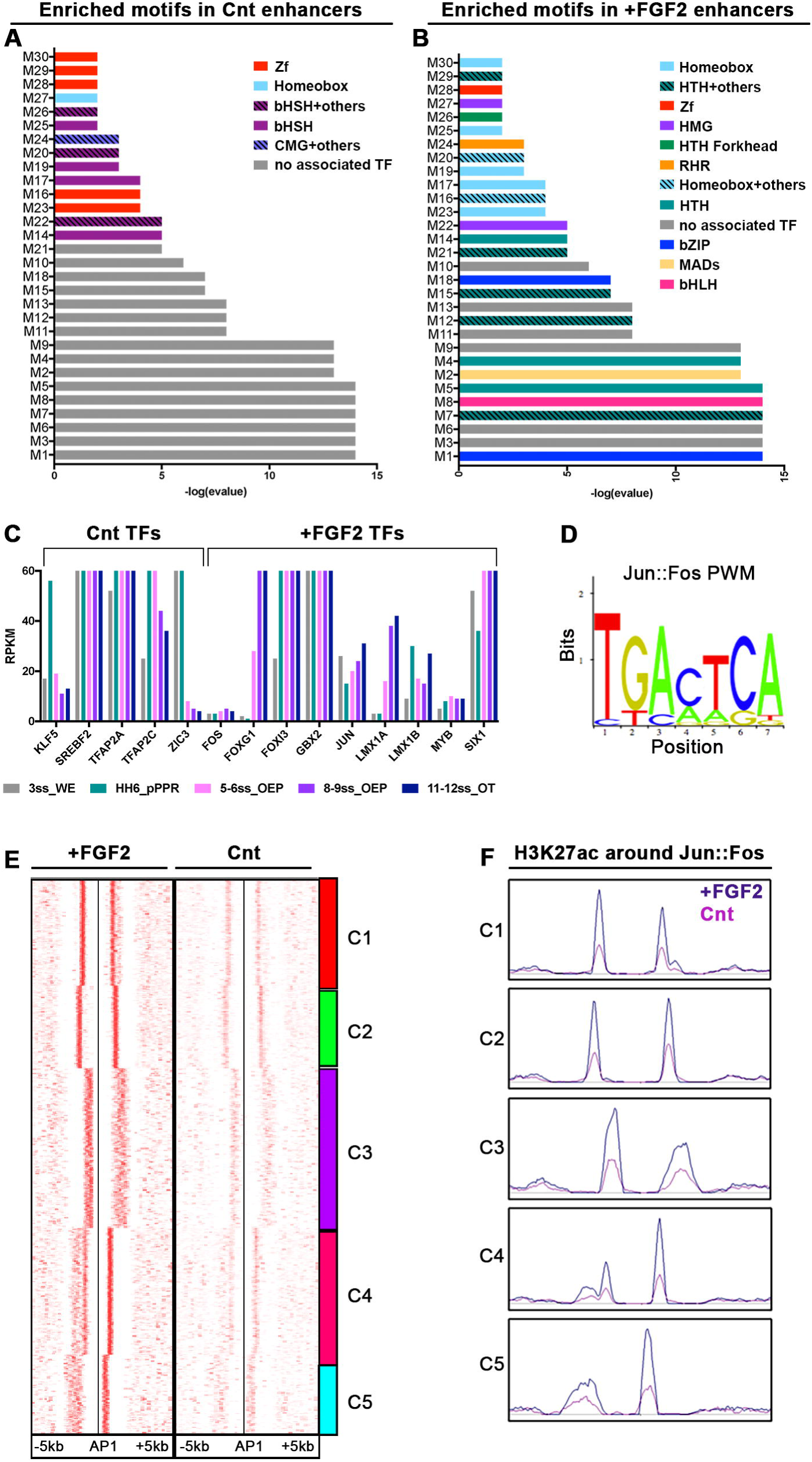
AP1 binding sites are associated with gain of H3K27ac. (A-B) Motif enrichment analysis using RSAT of enhancers active in control (Cnt; A) and FGF2 (B) treated sensory progenitors. Transcription factor families corresponding to the enriched motifs (M1-M30) are colour coded, -log(evalue) is plotted on the y axis. (C) Expression levels of transcription factors corresponding to enriched motifs in the 3ss whole embryo (grey), in 0ss posterior sensory progenitors (green), 5-6ss OEPs (pink), 8-9ss OEPs (violet) and the 11-12ss otic placode (blue) from (Chen et al., 2017). (D) Jun::Fos (Ap1) Position Weight Metrix (PWM) logo; note: the Ap1 motif is highly enriched in FGF2-treated samples. (E) Distribution of H3K27ac peaks surrounding AP1 motifs in FGF2-treated and control sensory progenitors; SeqMINER view of the heatmaps. (D) H3K27ac peak profiles around the Ap1 motif in FGF2 treated (violet) and control (pink) sens0ry progenitors in a window of -/+5kb.

**Figure 4.**
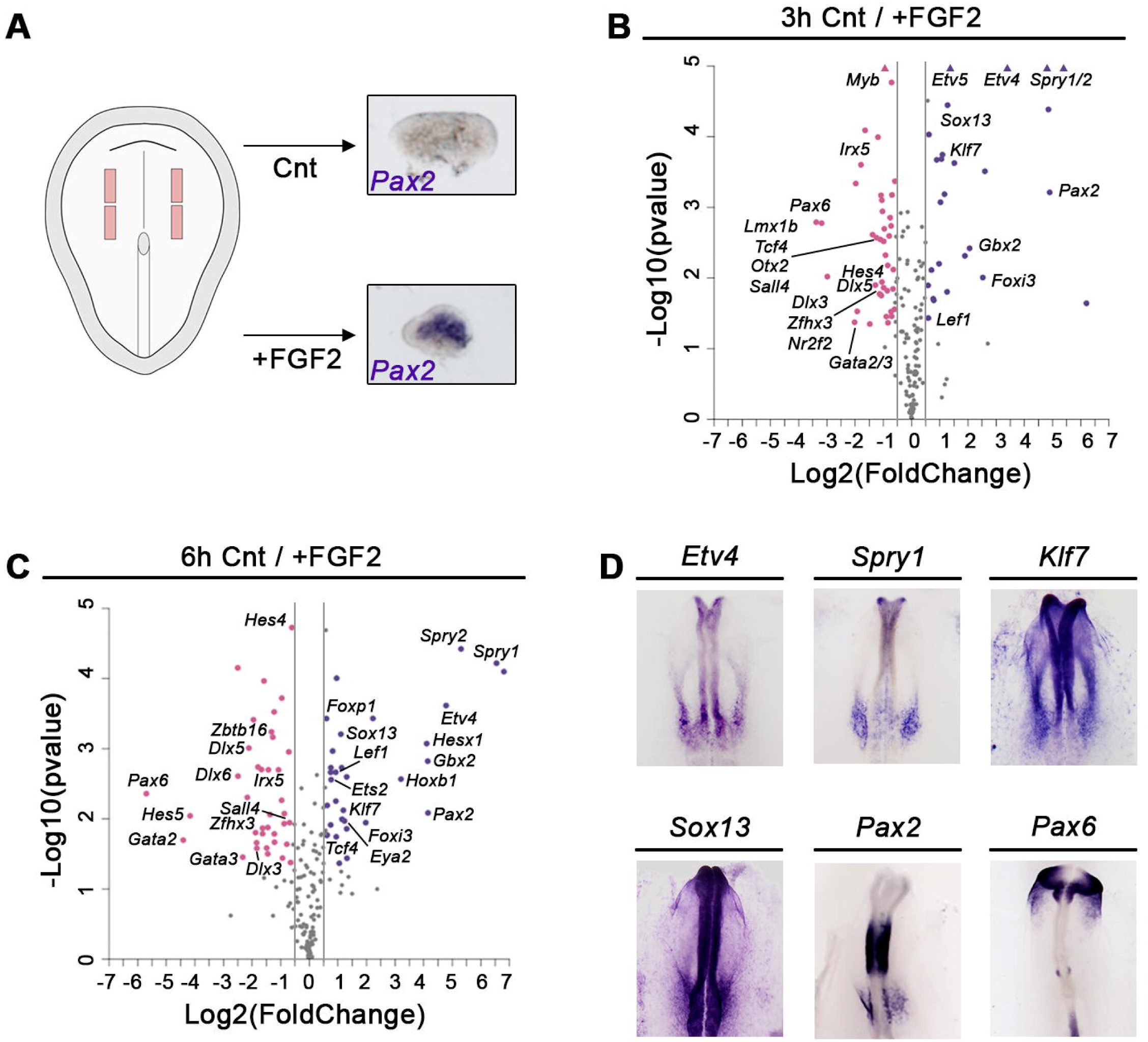
Enhancer associated transcripts are regulated by FGF signalling. (A) Sensory progenitor explants from HH6 chick embryos were cultured in the absence or presence of FGF2. After 6 hrs, the OEP marker *Pax2* is upregulated. (B) 3 hr FGF2 treatment promotes the expression of OEP transcripts, while repressing non-otic and late otic genes as determined by NanoString nCounter. A fold change of 1.5 or 0.25 (grey lines) and a p-value < 0.05 were used as threshold; transcripts not passing these thresholds are shown in grey and significantly up-and downregulated genes are shown in pink and violet, respectively. (C) After 6 hrs of FGF2 treatment sensory progenitors continue to express many 3hr-induced transcripts and upregulate new genes. A fold change of 1.5 or 0.25 (grey lines) and a p-value < 0.05 were used as threshold; transcripts not passing these thresholds are shown in grey and significantly up-and downregulated genes are shown in pink and violet, respectively. (D) Genes upregulated by FGF2 are expressed in OEPs, while FGF2-repressed genes are absent from OEPs (*Pax6*) as determined by in situ hybridisation.

To assess whether the genomic regions flanked by H3K27ac are indeed active OEP enhancers *in vivo*, we selected elements associated to known FGF targets or OEP genes, cloned them into reporter constructs to drive eGFP and assessed their activity in chick embryos. HH6 embryos were electroporated with enhancer-eGFP vectors together with a control vector driving RFP ubiquitously. There are four putative enhancers near *Spry1*, of which two were tested for *in vivo* activity. Spry1-E2 is conserved across vertebrate species as determined by DREiVE (Khan et al., 2012) (Figure 2A) and drives eGFP in otic progenitors from the 2-somite stage (ss) onwards (Figure 2B, C) recapitulating the normal expression pattern of *Spry1* (compare to Figure 4D), while E1 shows no activity (not shown). Two evolutionary conserved elements are associated with *Foxi3* (*Foxi3-E1* and *-E2*; Figure 2D), a transcription factor crucial for ear development (Birol et al., 2016; Khatri et al., 2014; Solomon et al., 2003); both drive reporter expression in two spatially and temporally distinct domains (Figure 2E, F; Supplementary Figures 5, 6). *Foxi3-E1* is active early in OEPs, but is shut down after the 5-6ss, while *Foxi3-E2* becomes active slightly later, when *Foxi3* is lost from the otic placode, and is confined to non-otic ectoderm. Thus, both enhancers recapitulate the dynamic expression of *Foxi3* in the head ectoderm (Khatri et al., 2014). In addition, enhancers associated with the otic transcription factor *Hesx1* and with the cytokine *Cxcl14* are also active in the otic territory (Supplementary Figure 6). Therefore, H3K27ac enrichment in response to FGF signalling identifies ear-specific enhancer regions.

**Figure 5.**
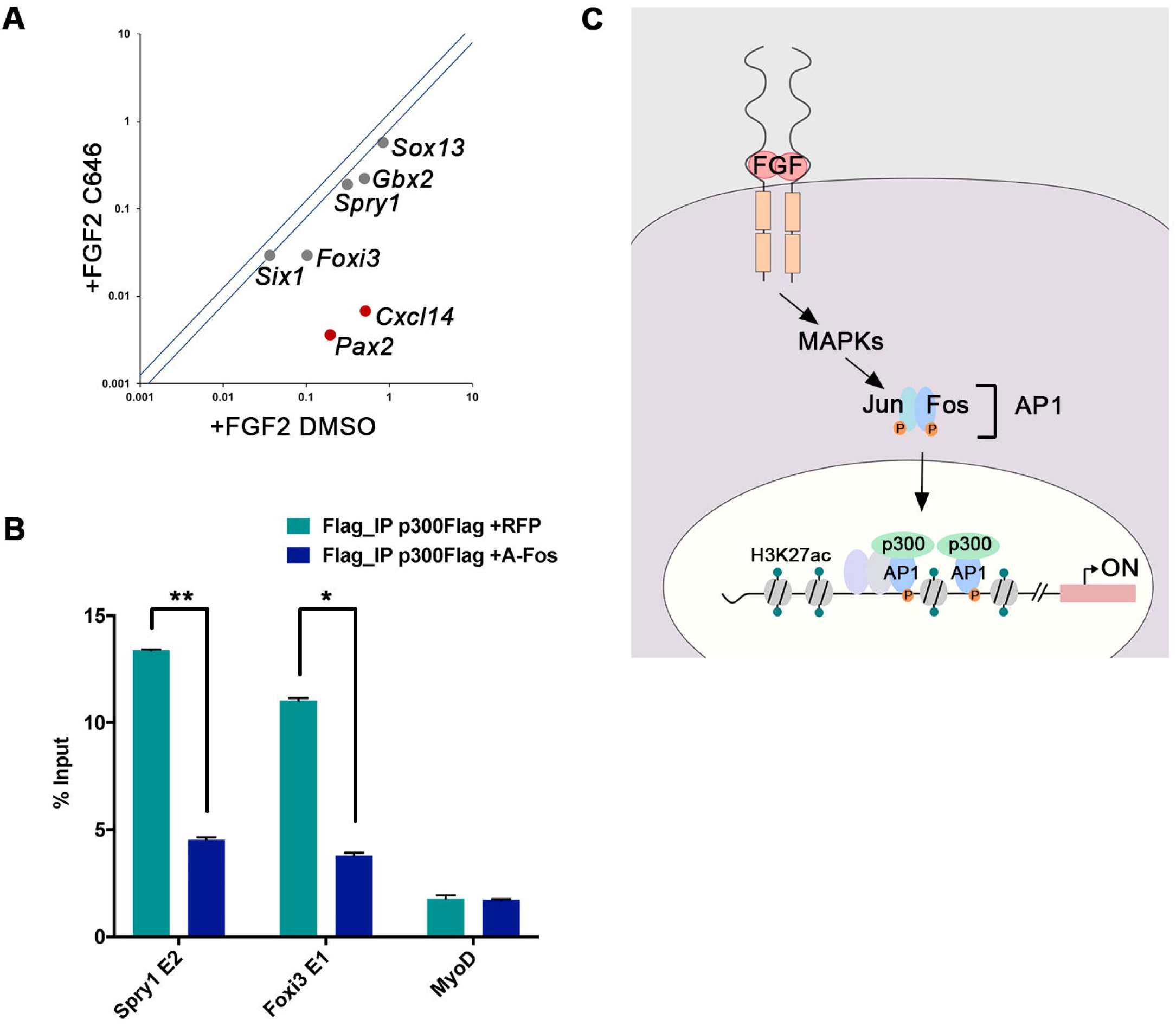
Ap1 recruits p300 to OEP enhancers. (A) De novo acetylation is required for OEP gene expression. Sensory progenitor explants from HH6 were cultured with FGF2 for 3 hrs in the presence of the p300 inhibitor C646 or vehicle control (DMSO). FGF2 fails to induce OEP genes when p300 activity is inhibited as assessed by qPCR. Red dots represent transcripts with significant difference, grey dots no significant changes. (B) p300 occupies OEP enhancers. Flag-tagged p300 together with a plasmid containing RFP (controls) or dominant negative A-Fos (experimental) was electroporated into future OEPs at HH6. After 6 hrs targeted otic placodes were dissected and ChIP was performed with anti-Flag antibodies. p300 occupies *Spry1* and *Foxi3* enhancers active in OEPs, but does not bind the enhancer for the muscle gene *MyoD*. In the presence of A-Fos p300 binding is abolished on both OEP enhancers. (C) A model for OEP gene activation in response to FGF signalling. See text for details.

### Unique transcription factor motifs define OEP enhancers

Enhancers serve as transcription factor hubs that integrate different transcriptional inputs to activate cell fate specific gene expression. To investigate whether a unique transcriptional signature might regulate OEP genes, we performed motif enrichment. Using RSAT matrix-scan we scanned for TF binding sites enriched in FGF2-treated versus control cells and vice versa using JASPAR and TRANSFAC libraries to identify known motifs. TFAP2 motifs (Figure 3A; Supplementary File1 - Cnt_M14_M17_M26) and motifs where the corresponding transcription factors are currently unknown (Supplementary File1) are highly overrepresented in control enhancers. In addition, they harbour Zic1/3, Srebf2 and Klf5 binding sites and the corresponding transcription factors *TFAP2a, -b, -c* and *-e, Klf5* and *Zic1/3* are expressed in sensory progenitors, while *Srebf2* seems to be expressed ubiquitously (Figure 3C; Supplementary Figure 7). Indeed, TFAP family members expressed in neural plate border cells control both neural crest and sensory progenitor fates (Hoffman, 2007; Knight et al., 2003; Li and Cornell, 2007; Qiao et al., 2012), while Zic transcription factors together with Pax3 discriminate between both fates (Hong and Saint-Jeannet, 2007).

In contrast, OEP associated enhancers show a different motif signature. Motifs corresponding to homeobox and winged-helix/forkhead domain transcription factors are highly enriched (Supplementary File2 FGF2_M5_12_15_21_16_20_29), as are binding sites for basic-loop-helix factors (Supplementary File2 FGF2_M8), for the transcription factor Myb (Supplementary File2 FGF2_M7) and the FGF signalling mediators of the Jun and Fos families (Figure 3B; Supplementary File2 FGF2_M18). Many of the corresponding transcription factors are expressed in OEPs and required for their specification including *Dlx5/6/3, Gbx2, Lmx1a* and *Six1* and the forkhead factor *Foxi3* (Birol et al., 2016; Brugmann et al., 2004; Chen et al., 2017; Christophorou et al., 2009; Kaji and Artinger, 2004; Khatri et al., 2014; Luo et al., 2001; McLarren et al., 2003; Sato et al., 2010), while others play a role at later stages of ear development (e.g. *Foxg1*) (Hwang et al., 2009) and yet others are expressed in OEPs, but their function has not been investigated (*FoxP1, FoxC2, FoxK2*; (Chen et al., 2017). Thus, a unique signature of transcription factor binding sites defines OEP enhancers and identifies new potential regulators of OEP identity.

The Fos/Jun transcription factors are mediators of FGF signalling and form homo-or heterodimers (AP1) to activate downstream targets. The AP1 motif was differentially associated with FGF2-induced elements (Figure 3B; Supplementary File1 – FGF2_M18) and is indeed present in the enhancers tested *in vivo* (Figure 2A, D; Supplementary Figure 5C). This finding suggests a major difference in enhancer activation during OEP induction can be attributed to the FGF pathway. Examining the distribution of H3K27Ac within 5kb surrounding AP1 motifs reveals a strong correlation in FGF-treated cells, while the same regions are depleted in controls (Figure 3E, F). Clustering and peak density plots identifies five different clusters (C1-C5) that differ in the location of H3K27ac marks with respect to AP1 motifs (Figure 3F). Functional annotation of these clusters shows enrichment of terms like ear morphogenesis, inner ear development, sense organ development and, importantly, MAPK signalling (Supplementary Figure 8). Together, these data define a unique set of transcription factor binding sites over-represented in FGF activated enhancers when compared to control enhancers. The corresponding transcription factors are known to control otic identity, while others represent new putative upstream regulators.

### Genes associated to FGF-activated enhancers are FGF targets in OEPs

The above experiments identified several thousand enhancers that are rapidly activated in response to FGF signalling. However, previous studies identified only few immediate FGF targets (Anwar et al., 2017) or few genes induced after 24hrs’ exposure to FGF (Yang et al., 2013a). We therefore investigated whether genes associated with FGF-activated enhancers are indeed regulated by this pathway. We have recently characterised the transcriptional profile of OEPs (Chen et al., 2017) and based on these data designed a NanoString probe set with 216 genes containing 70 ear specific factors, as well as transcripts normally expressed in progenitors for other sense organs, cranial ganglia, neural and neural crest cells. Sensory progenitors from HH6 chick embryos were cultured in the absence or presence of FGF2 for 3 and 6 hours and changes in gene expression were evaluated by NanoString (Figure 4A). After 3 hrs known FGF targets (*Etv4/5, Spry1/2*) are strongly up-regulated together with early OEP markers (e.g. *Foxi3, Gbx2, Pax2, Sox13* and *Klf7;* in total 16 otic TFs), while genes normally expressed in other cell types (e.g. *Pax6, Otx2, Msx1, Id2, -4*) and some late otic genes (*Sall4, Zfhx3, Fez1, Lmx1b, Myb*) are repressed (Figure 4B, D; Supplementary File 3). After 6 hrs of FGF exposure, the expression of most early induced transcripts is maintained and a few new factors become upregulated (e.g. *Eya2, N-Myc, Tcf7L2*; Figure 4C; Supplementary File 4). These changes in gene expression correlate well with enhancer activation by FGF signalling: most upregulated transcripts are associated with an FGF-induced gain of H3K27ac (Supplementary Figure 3, 4). Together, these experiments suggest that FGF signalling initiates OEP induction by directly activating enhancers of ear specific genes, and that this activity maybe mediated by AP1 together with other transcription factors.

### AP1 recruits the histone acetyl-transferase p300 to ear-specific enhancers

What is the mechanism by which FGF and AP1 activate OEP enhancer regions? Acetylation of H3K27 is mediated by histone acetyl-transferases (HAT) like p300. To assess whether p300 occupies otic enhancers, we selected two experimentally verified enhancers, *Spry1-E1* and *Foxi3-E1*, with a significant gain of H3K27ac in response to FGF activation. We electroporated p300-Flag into ear progenitors and performed anti-Flag ChIP-qPCR from targeted tissue. Indeed, p300 is bound to both enhancers, but is absent from the enhancer of the muscle gene *MyoD* (Figure 5B). To test whether *de novo* acetylation is required for OEP gene expression we treated sensory progenitors with FGF2 in the presence or absence of the p300 inhibitor C646 (Bowers et al., 2010; Zhang et al., 2013) and assessed OEP markers after 3 hrs by qRT-PCR. When p300 is inhibited the expression of the FGF-dependent OEP transcripts *Pax2* and *Cxcl14* is no longer induced. In addition, there is a non-significant reduction of *Spry*1, *Gbx2, Foxi3* and *Sox13*, while *Six1* does not change in line with its FGF independence (Figure 5A). Using the general HAT inhibitor anacardic acid has the same effect (data not shown). Therefore, *de novo* acetylation of p300 targets is required for OEP gene expression.

AP1 is known to interact with p300 (Crish and Eckert, 2008), raising the possibility that AP1 recruits p300 to OEP enhancers to facilitate histone acetylation. To test this, we made use of a dominant negative form of Fos (acidic-Fos: a-Fos; (Biddie et al., 2011), a truncated version of Fos that maintains its interaction with Jun, but cannot bind DNA. HH6 chick embryos were electroporated with p300-Flag together with a-Fos or RFP (control) constructs; targeted otic progenitors were dissected and processed for ChIP-qPCR using anti-flag antibodies. While p300 binding to *Spry1-E1* and *Foxi3-E1* is observed in controls, this is inhibited in the presence of a-Fos (Figure 5B). No binding is observed at the MyoD enhancer, a transcript not expressed in ear progenitors. These findings suggest that in response to FGF signalling, AP1 recruits the HAT p300, which in turn increases acetylation of H3K27 at ear-specific enhancers and as a result rapidly induces their expression (Figure 5C).

## DISCUSSION

### FGF signalling mediates enhancer activation in otic-epibranchial progenitors

In the embryo, cell fate choices are mediated by temporally and spatially controlled signals that activate distinct transcriptional programmes in responding cells. The key elements that integrate this information are non-coding, cis-regulatory regions that control cell type specific gene expression, which in turn implement fate decisions (Banerji et al., 1981; Beagrie and Pombo, 2016; Pennacchio et al., 2013). Enhancer elements are characterised by specific epigenetic signatures, however how transient signalling cues during development affect the chromatin landscape is not well understood. Active enhancers are nucleosome-free regions flanked by characteristic histone modifications including acetylation of histone 3 at lysine 27 (Creyghton et al., 2010; Kharchenko et al., 2011; Rada-Iglesias et al., 2012; Rada-Iglesias et al., 2011; Zentner et al., 2011). Here we study the changes in H3K27ac as sensory progenitors acquire OEP fate in response to FGF signalling. We find that FGF rapidly induces dynamic changes at thousands of genomic regions with a marked gain of H3K27 acetylation near OEP genes. We show that these act as enhancer elements that drive OEP-specific gene expression *in vivo* and enhancer associated transcripts are indeed rapidly upregulated by FGF. In contrast, genomic regions surrounding genes normally absent in OEPs lose H3K27 acetylation, and these transcripts are repressed by FGF. Thus, we have identified thousands of FGF-activated and - repressed regulatory elements during OEP induction providing a rich resource to explore the gene regulatory network that controls ear development.

### AP1 mediates p300 recruitment to FGF activated enhancers

FGF signalling activates gene expression in responding cells within minutes, and our studies provide a molecular mechanism underlying this response. In ear progenitors, FGF is mediated by the MAP-kinase pathway (Yang et al., 2013a) leading to phosphorylation of the transcription factors Jun and Fos, which work as a heterodimer AP1 to activate gene expression (Ornitz and Itoh, 2015; Tsang and Dawid, 2004; Yang et al., 2003, 2013b). The AP1 binding motif is strongly enriched in FGF-activated enhancers, and we show that AP1 is required to recruit the histone acetyltransferase p300 to ear enhancers. Indeed, AP1 and p300 physically interact (Crish and Eckert, 2008), and inhibition of p300 prevents induction of FGF response genes in sensory progenitors. This supports the idea that p300 activity and subsequent increased histone acetylation is required to activate OEP genes. A similar mechanism has recently been reported in HeLa cells (O’Donnell et al., 2008). Here, in response to MAP kinase signalling, Elk is phosphorylated and together with p300 changes the histone acetylation state, which in turn allows the recruitment of other transcription factors and subsequent gene activation. It is therefore possible that increased acetylation in OEPs enhances chromatin accessibility allowing other otic transcription factors to access target enhancers, where they may cooperate with AP1 to activate target transcription. Motif enrichment analysis of FGF-induced ear enhancers suggests that homeobox and winged-helix/forkhead proteins are possible cooperators. Of those, the homebox genes *Gbx2, Six1* and *Lmx1a* and the forkhead gene *Foxi3* are indeed expressed in OEPs and are required for their specification and/or vesicle formation (Birol et al., 2016; Brugmann et al., 2004; Chen et al., 2017; Christophorou et al., 2009; Kaji and Artinger, 2004; Khatri et al., 2014; Luo et al., 2001; McLarren et al., 2003; Sato et al., 2010). In addition, we have recently shown that Myb recruits the histone demethylase Lsd1 to OEP gene promoters to maintain their expression after initial specification (Ahmed and Streit, 2018), while our current results suggest that Myb may also cooperate with AP1 at OEP enhancer regions.

In addition, AP1 is known to influence chromatin accessibility using different mechanisms and may act as a pioneer factor to mediate chromatin opening. In mouse embryonic fibroblasts, AP1 can bind to nucleosome-occupied regulatory elements together with cell type specific transcription factors (Vierbuchen et al., 2017). Together, they recruit the SWI/SNF complex BAF, which repositions nucleosomes to increase enhancer accessibility. Likewise, AP1 is required for glucocorticoid induced transcription by priming glucocorticoid receptor target sites and maintaining their accessibility before the receptor can be recruited (Biddie et al., 2011). Whether similar mechanisms also operate in OEPs in response to FGF signalling remains to be uncovered.

FGF signalling not only promotes OEP fate, but also shuts down alternative differentiation programmes. For example, we have previously shown that the lens programme is inhibited in OEPs by FGFs with the lens marker *Pax6* being a direct target (Anwar et al., 2017; Bailey et al., 2006). Here we observe a significant decrease of H3K27ac in the Pax6 locus and a complementary increase of H3K27me3. Recent evidence suggests that AP1 may mediate transcriptional repression by recruiting histone deacetylases (Miotto et al., 2006; Mittelstadt and Patel, 2012), which subsequently allows H3K27 methylation. Interestingly, in the neural tube FGF signalling promotes compaction of the Pax6 locus mediated by polycomb repressive complexes (Patel et al., 2013; Semprich et al., 2019), while inhibition of Erk1/2 results in dissociation of this complex from the Pax6 TSS. It is therefore possible that in OEPs increased H3K27me3 deposition at the Pax6 locus allows recruitment of the polycomb repressor complex to repress *Pax6*, a key gene for lens formation. Thus, AP1 function depends on a fine balance between activator and repressor complexes, presumably controlled by the availability of different, enhancer specific co-factors.

### OEP induction: new transcription factors downstream of FGF signalling

It is well established that FGF signalling activates the inner ear programme by inducing otic-epibranchial progenitors from a pool of multipotent precursors. While continued FGF activation promotes epibranchial identity, Notch together with Wnt signalling establishes the otic placode (Freter et al., 2008; Ohyama et al., 2006) (for review (Chen and Streit, 2013; Groves and Fekete, 2012; Ladher, 2017; Ohyama et al., 2007; Whitfield, 2015). Here we identify new FGF-response genes and their associated enhancers. Using network inference approaches we have recently proposed that FGFs initially enhances the transcription of genes already expressed in sensory progenitors, which then act in a positive feedback loop to increase their own expression (Anwar et al., 2017). Downstream of this circuit, *Pax2* and other transcription factors are activated to regulate OEP identity while repressing alternative fates. Here, we show that within 3 hrs FGF leads to the activation of 20 transcription factors, many of which are expressed in OEPs, but absent from other placodes. These include the known targets *Etv4* and *-5* and the FGF antagonists *Spry-1* and *-2*, as well as many new FGF-responsive factors like *Sox13, Tead3* and *Klf7*. Transcription factor binding motifs for these factors are present in ear enhancers suggesting that they may also play a key role in the genetic programme controlling early ear induction. Indeed, in *Xenopus* and mouse, the transcription factor *Klf7* is expressed in the otic placode (Gao et al., 2015; Laub et al., 2001). Klf7 has been implicated in stem cell renewal, as well as cell cycle exit of neuronal precursors (Jeon et al., 2016; Kajimura et al., 2007; Laub et al., 2006; Laub et al., 2005), however its function in ear development remains unknown.

In summary, during the induction of otic-epibranchial progenitors, FGF signalling rapidly activates downstream target genes by increasing H3K27 acetylation at enhancer regions. FGF activity is mediated by the Jun/Fos heterodimer AP1, which recruits the acetyltransferase p300 to OEP enhancers. The genome-wide identification of FGF-responsive OEP enhancer regions forms the basis to reconstruct the otic induction network and may shed light on other FGF-mediated processes in the ear.

## METHODS

### Embryo manipulation and explant culture

Experiments on chick embryos prior to E10 do not require a home office license or institutional approval and were carried out according to the institutional guidelines. Fertilized chicken eggs were obtained from Winter Farm (Herts, UK) and incubated in a humidified incubator at 38°C until reaching the desired stage (Hamburger and Hamilton, 1951). For electroporation, embryos were cultured using the filter culture method (Chapman et al., 2001). To isolate sensory progenitor explants, embryos were collected in Tyrode’s saline and progenitors were dissected using fine needles, and cultured in DMEM supplemented with FGF2 (0.25ng/μl; R&D), DMSO or C646 (10μM; Calbiochem) for 3 and 6 hours.

### Enhancer cloning and electroporation

Putative enhancer elements were amplified from chick genomic DNA and inserted into pTK vector containing eGFP reporter and a TK minimal promoter (Kondoh and Uchikawa, 2008). Cloning primers are listed in Supplementary Table1. Flagged tagged p300 vector was kindly provided by Michael O. Hottiger (Hasan et al., 2001). For electroporation, embryos on filter paper were transferred into an electroporation chamber containing a 2×2mm platinum electrode. DNA (3 μg/μl reporter DNA pTK eGPF and 1.5 μg/μl pCAB RFP, 0.1% fast green; 1 μg/μl p300-Flag and 1.5 μg/μl pCAB RFP, 0.1% fast green) was injected by air pressure between the vitelline membrane and embryonic ectoderm, a platinum electrode (2×1 mm) was placed above the target area and 5 pulses of 4V for 50ms with 750ms intervals were applied using Intracel TSS20 OVODYNE pulse generator. Embryos were cultured overnight and imaged using an inverted fluorescent microscope.

### Quantitative RT-PCR and NanoString nCounter analysis

RNA from cultured sensory progenitor explants treated with FGF2 (0.25ng/μl; R&D), DMSO or C646 (10μM; Calbiochem) for 3 hrs was isolated using the RNAqueous-Micro Kit (Ambion) and reverse transcribed. Primers for target genes were designed with PrimerQuest (IDT). qPCR was performed in technical triplicates using Rotor-Gene Q (Qiagen) with SYBR green master mix (Roche). The ΔΔCt method was used to calculate the fold change, which was expressed as FC=2∼(-ΔΔCt) (Livak and Schmittgen, 2001). Gapdh and Sdha were used as reference genes to calculate the fold change. Two biological replicates were performed and the P-value generated from a two-tailed Student’s t-test was used to evaluate statistical significance.

For Nanostring nCounter analysis, eight to ten explants per condition were lysed in 5μl lysis buffer (Ambion) and analyzed by nCounter® Analysis System (Life Sciences) using a customized probe set of 216 genes (Chen et al., 2017). Total RNA was hybridized with capture and reporter probes at 65 °C overnight. Target/probe complexes were washed and immobilized according to the nCounter Gene Expression Assay Manual, and data were collected by the nCounter Digital Analyzer. Experiments were repeated on three independent occasions and data were analyzed following company instructions. A cut-off of fold change >= 1.25 and <=0.75 was used to identify upregulated and downregulated genes, respectively, in combination with a p-value <=0.05 (unpaired t-test).

### *In situ* hybridization

The following chick plasmids were used to generate Digoxigenin–labeled antisense probes: Etv4 (a kind gift from M. Bronner) Spry1 obtained from Life Technologies Klf7 `ChEST376015`; Sox13 ChEST437d11; Pax2 (a kind gift from M. Golding) and Pax6 (a kind gift from A. Bang). RNA probes were synthesized with T7, T3 or SP6 RNA polymerase (Roche). Whole mount or explant *in situ* hybridization was performed as described previously (Streit and Stern, 2001).

### Chromatin immunoprecipitation (ChIP)

Around 100 posterior sensory progenitors from HH6 were cultured for 6 hours in the absence or presence of FGF2. Explants were dissociated in 0.5ml of Nuclei Extraction Buffer (NEB; 0.5% NP-40, 0.25% Triton X-100, 10mM Tris-HCl pH 7.5, 3mM CaCl_2_, 0.25M Sucrose, 1mM DTT, 0.2mM PMSF, 1X Protease Inhibitor) by homogenisation and crosslinked using 1% formaldehyde for 9 min at room temperature. 0.125M glycine final concentration was used to stop fixation. Samples were pelleted and stored at −80°C. 100μl of Dynal magnetic beads (Protein A, Novex Life Technologies) were incubated with 1ml of blocking buffer (1X PBS with 0.5% BSA) for 2 min, resuspended in 250μl of blocking buffer and respective antibodies were added (1μg/μl Rabbit anti-IgG Millipore – CA92590; 1μg/μl Rabbit anti-H3K27ac Abcam – ab4729; 2.5μg Rabbit anti-H3K27me3, Cell signaling – C36B11; 3μg anti-Flag, Sigma – F3165). Antibody and beads were incubated overnight rotating at 4°C, and excess of antibody was removed by washes in blocking buffer. Crossed-linked pellets were defrosted, washed in Nuclei Extraction Buffer and nuclei were extracted by homogenisation. Samples were lysed in 1% SDS, 50mM Tris-HCl (pH 8), 10mM EDTA supplemented with 20μl Protease Inhibitor (7x PI; Roche) for 1 hour. Prior to sonication 280μl of ChIP dilution buffer (0.01% SDS; 16.7mM Tris-HCl pH 8; 1.2mM EDTA; 167mM NaCl; supplemented with PMSF, DTT and PI) was added, samples were sonicated to generate fragments of 200-600bp using 12 cycles of 15sec on and 30sec off at 40% amplitude (SONICS, Vibra Cell™). The resulting chromatin was diluted 4x and divided across different samples, 1/10^th^ of sonicated chromatin was kept frozen as input control. Beads were washed 9 times in RIPA buffer containing 50mM Hepes-KOH (pH 8), 500mM LiCl, 1mM EDTA, 1% NP-40, 1.7% Na-deoxycholate supplemented with PI. A final wash was performed using 1x TE, 50mM NaCl and the chromatin was eluted in 50mM Tris-HCl (pH 8), 10mM EDTA and 1% SDS. Reverse-crosslinking was carried out at 65°C overnight. Chromatin was diluted using 1xTE and incubated at 37°C for one hour with RNAse A 0.4mg/μl, followed by proteinase K digestion for one hour at 55°C. Chromatin was purified using phenol: chloroform: isoamyl alcohol and resuspended in 60μl of water. ChIPed chromatin was processed for next generation sequencing or analyzed by qPCR. For qPCR Ct values for each precipitation were extracted and normalized to input chromatin, primer sequences are listed in Supplementary Table 2. For next generation sequencing chromatin was amplified with a step of linear amplification following the protocol described in (Adli and Bernstein, 2011) Library preparation was performed at the UCL Genomics, Institute of Child Health following the protocol used for nano-ChIP-seq (Adli and Bernstein, 2011) with the exception that only 12 cycles were used in the PCR amplification.

### ChIP-seq data analysis

Sequence quality was assessed using FastQC (Andrews, 2010). During each ChIP-seq experiment, an amplification step was carried out that is reported to produce mismatches at the first 9bp due to random priming (Adli and Bernstein, 2011). Therefore, as the sequence quality was poor these 9bp were trimmed. Further trimming was performed to improve alignment if sequence quality from the 3’ end was poor. Reads were aligned to the chick genome Galgal4.71 using Novoalign (Novocraft 2.08.01, http://www.novocraft.com/products/novoalign/) and uniquely aligned sequences were used for peak calling using Homer (Heinz et al., 2010) and MACS2 (Zhang et al., 2008). For Homer, a fold change of 1.5 relative to input and a False Discovery Rate (FDR) of 0.01 were used. For MACS2, a FDR of 0.05 as suggested in the MACS manual to obtain broad peaks (characteristic of histone peaks) and a default p-value of 1e-5 were used. Genome wide analysis was performed on Homer called peaks. Following peak-calling, putative enhancers were identified in the following way: regions of up to 3 kb flanked by H3K27ac and devoid of H3K27me3 peaks were identified and assigned to the nearest gene using gene annotations from Ensembl (Galgal4.71) and refGene (Nov. 2011 ICGSC Gallus_gallus-4.0/galGal4). Read distributions around transcription start site (TSS) or the centre of a putative enhancer was plotted using ’annotatePeaks’. Putative enhancers for +FGF2-treated and control samples were compared to find common and unique putative enhancers using the R package ChIPpeakAnno (Zhu et al., 2010). Putative enhancers in +FGF2 and -FGF2 were considered to be overlapping and common if they had a 0 bp gap between them, otherwise they were considered to be unique to the respective condition. Genes associated with unique +FGF2 and -FGF2 putative enhancers were subjected to Gene Ontology (GO) analysis using DAVID (DAVID Bioinformatics Resources 6.7) (Huang et al., 2009a, b). All ChIP-seq data were viewed in the IGB browser (Nicol et al., 2009).

### Transcription factor binding site analysis

Transcription factor binding site analysis was carried out using RSAT peak-motifs (Thomas-Chollier et al., 2012a; Thomas-Chollier et al., 2012b) and JASPAR (Sandelin et al., 2004) and TRANSFAC (Matys et al., 2006) libraries. To obtain significant binding sites, an e-value <= 0.05 was considered.

### Conservation

Multiple alignments between 21 amniotes were obtained from Ensemble PECAN (Paten et al., 2008). Additionally, DREiVe (Khan et al., 2012), a motif-discovery algorithm was used to identify putative regulatory regions 300 KB upstream and downstream of selected FGF-response genes. It reports regulatory regions as clusters of short conserved motifs of 8 bp in a 300 bp window. DREiVe does not depend on sequence alignment, is able to identify re-arrangements of motifs within regulatory elements and does not require prior information of transcription factor binding sites. Regions that were conserved in 7 out of 9 species (human, horse, cow, rabbit, mouse, opossum, platypus, chick and lizard) were considered as putative enhancers. These tracks were then loaded into IGB browser to view with Chip-seq data.

## ACKNOWLEDGEMENTS

The authors would like to thank Ms Ewa Kolano for excellent technical support, Jingchen Chen and Ramya Ranganathan for help with tissue collection and data analysis, and Claudio D. Stern, Nicolas Luscombe and Alvaro Rada-Eglesias for critical reading of the manuscript and the Streit group for helpful discussions. This work was funded by research grants from the BBSRC (BB/M006964/1), NIH Deafness Research UK (513:KCL:AS) and the National Institute on Deafness and Other Communication Disorders (DC011577).

## AUTHOR CONTRIBUTIONS

A.S. designed the experiments together with M.T.; M.T. conducted most experiments and analysed the data together with A.S.; M.A. performed all bioinformatics analysis; M. Ahmed performed ChIP-qPCR in figure 5. M.T. and M. A. prepared the figures. A.S wrote the manuscript.

