## Supplementary material for "FGF signalling regulates enhancer activation during ear progenitor induction"

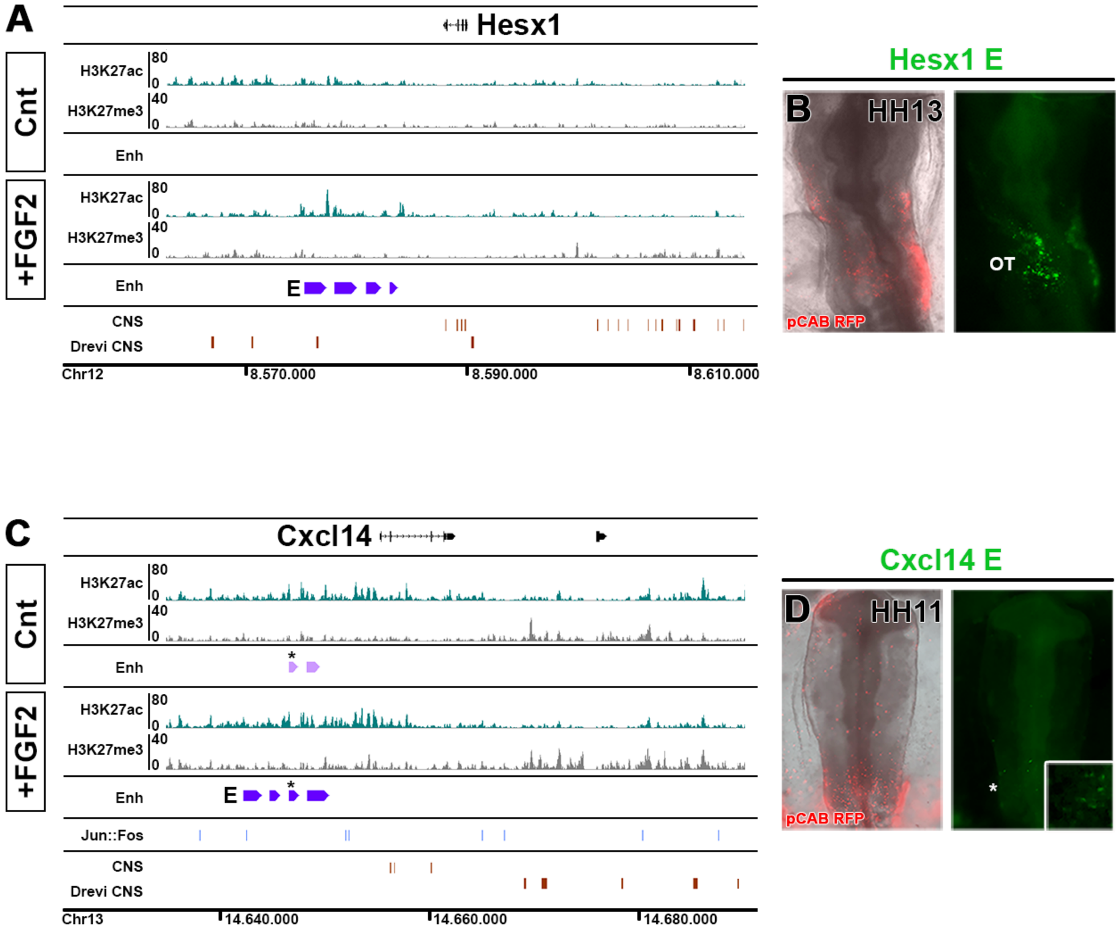


**Supplementary Figure 1. Hesx1 and Cxcl14 enhancers.**

IGB browser view of Hesx1 (A) and Cxcl14 (C) enhancers. ChIP-identified enhancers are in violet for +FGF2 and pink for control samples (* marks common enhancers between Cnt and +FGF2); H3K27ac track is shown in green and H3K27me3 in grey. Jun::Fos putative binding sites are shown for Cxcl14 locus (blue); conserved non-coding sequences (CNS) are in red. Characterization of *in vivo* activity of Hesx1 E (green channel) shows that it is active in the otic placode (OT) at HH13 (B). The Cctive xcl14 enhancer element is ain few cells in the neural tube and in the ectoderm surrounding the otic placode (magnified inset) (D).

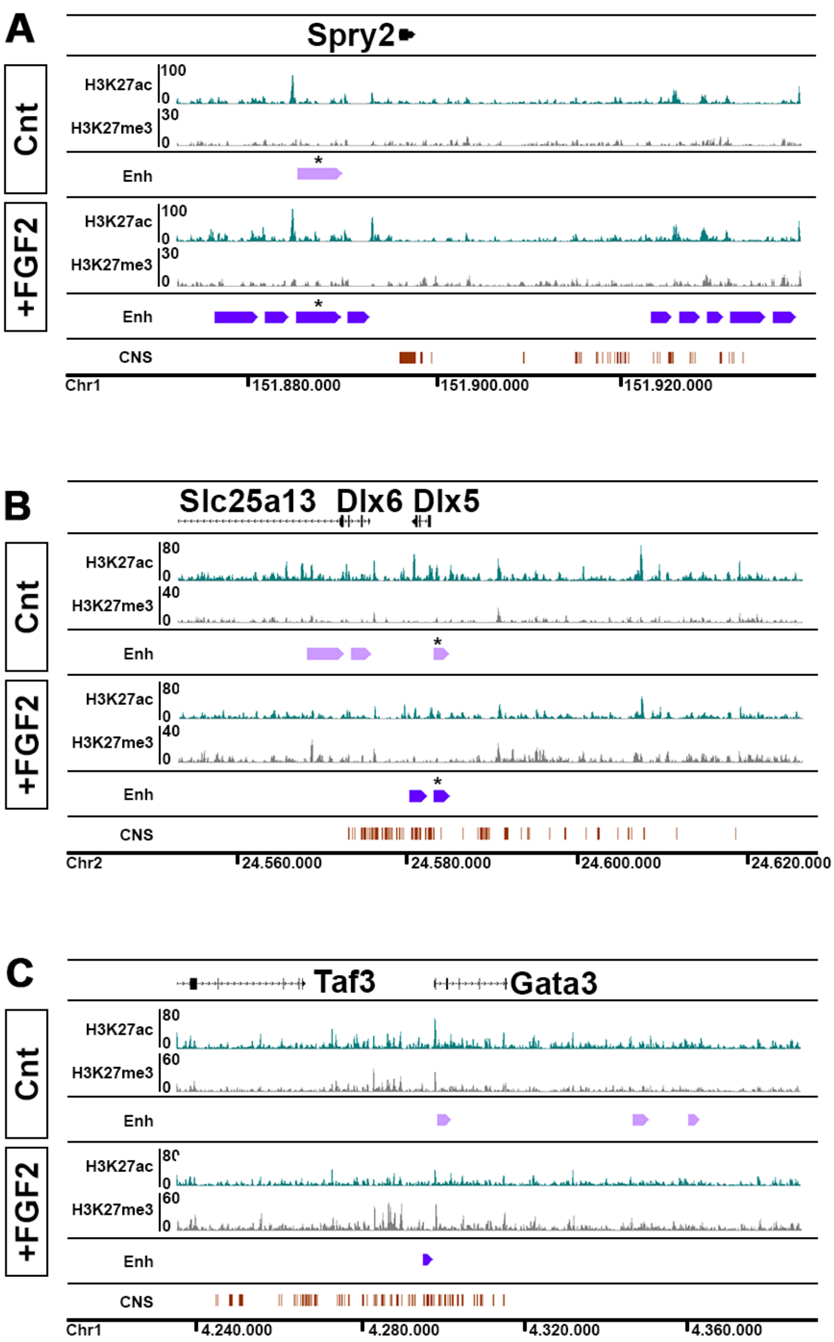


**Supplementary Figure 2. Genomic regions surrounding Spry2, Dlx5/6 and Gata3.**

IGB browser view of the Spry2 (A), Dlx5/6 (B) and Gata3 (C) locus. ChIP-identified enhancers are in violet for +FGF2 and pink for control samples (* marks common enhancers between Cnt and +FGF2); H3K27ac track is shown in green and H3K27me3 in grey. Conserved non-coding sequences (CNS) are in red. FGF2 induction increases H3K27ac around Spry2 (A) while there is a decrease in H3K27ac and a gain in H3K27me3 in Dlx5/6 (B) and Gata3 (C), which are genes negatively regulated by FGF signalling.

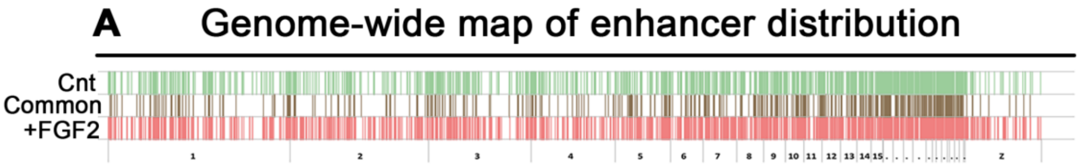


**B**

FGF2-repressed genes associated with enhancers

HES1
BMP2
TBX3
TBX2
JARID2
NMRAL1
BCL11A
TLE4
MXI1
CBFA2T2
DNMT3B
N2C1

**C**

FGF2-activated genes associated with enhancers

CXCL14
SPRY1
SPRY2
HESX1
EPHA4
FOXI3
BMP4
FOXP1
CHD7
CHD2

**Supplementary Figure 3. Genome-wide map of enhancers identified in control and +FGF2 treated sensory progenitors.**

(A) A genome-wide view of the location of common and unique enhancers in +FGF2 and control samples. (B) List of genes significantly downregulated by FGF2 with an associated proximal enhancer determined by ChIP-seq in control sensory progenitors. (C) List of genes significantly upregulated by FGF2 with an associated proximal enhancer determined by ChIP-seq in FGF2-treated sensory progenitors.

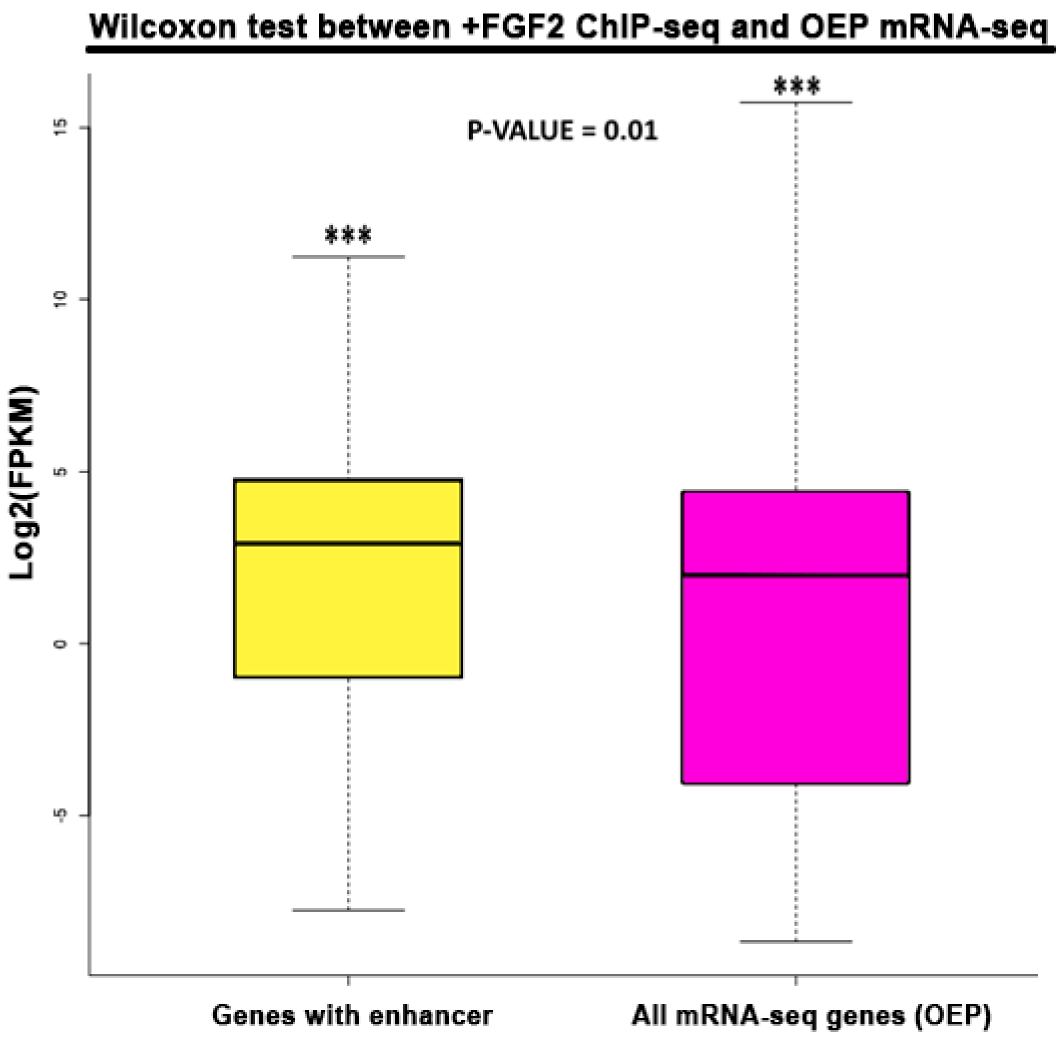


**Supplementary Figure 4. Wilcoxon test reveals a significant correlation between +FGF2 ChIP-seq and OEP mRNA-seq.**

Putative enhancers in FGF2 treated sensory progenitors were defined as maximum 3 kb genomic regions flanked by H3K27ac peaks and devoid of H3K27me3 peaks. These were annotated to the nearest gene. mRNA-seq for OEPs were retrieved from Chen and colleagues (Chen et al., 2017). A Wilcoxon test was carried out to test if the mean FPKM of genes with putative enhancers in FGF2 treated sensory progenitors is greater than the mean FPKM of all OEP genes. This analysis shows that indeed enhancer associated genes have high expression levels in OEPs with a significant p-value of 0.01.

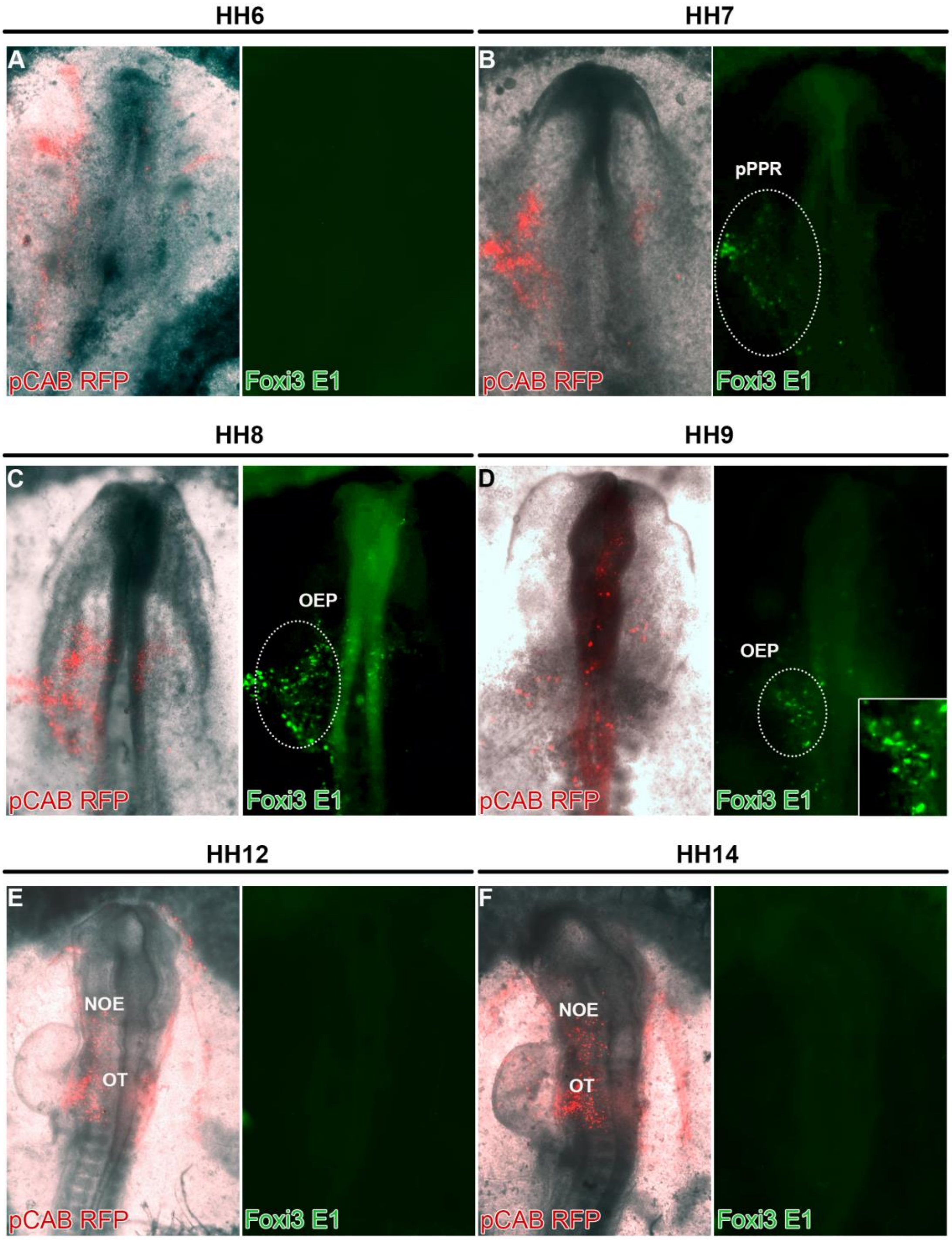


**Supplementary Figure 5. *In vivo* activity of the Foxi3-E1 enhancer.**
Foxi3-E1 driving eGPF and b-actin driving RFP were coelectroporated into chick embryos at primitive streak stages, and enhancer activity monitored from 0-1ss onwards. A few Foxi3-E1 GFP+ cells were first identified at HH7 in posterior sensory progenitors (pPPR) (A, B); between HH8-9 the enhancer shows broad activity in OEPs (C, D) and it becomes inactive around HH12 (E, F). NOE: non-otic ectoderm, pPPR: posterior Pre-Placodal Region, OEP: Otic-epibranchial Placode, Ot: otic placode.

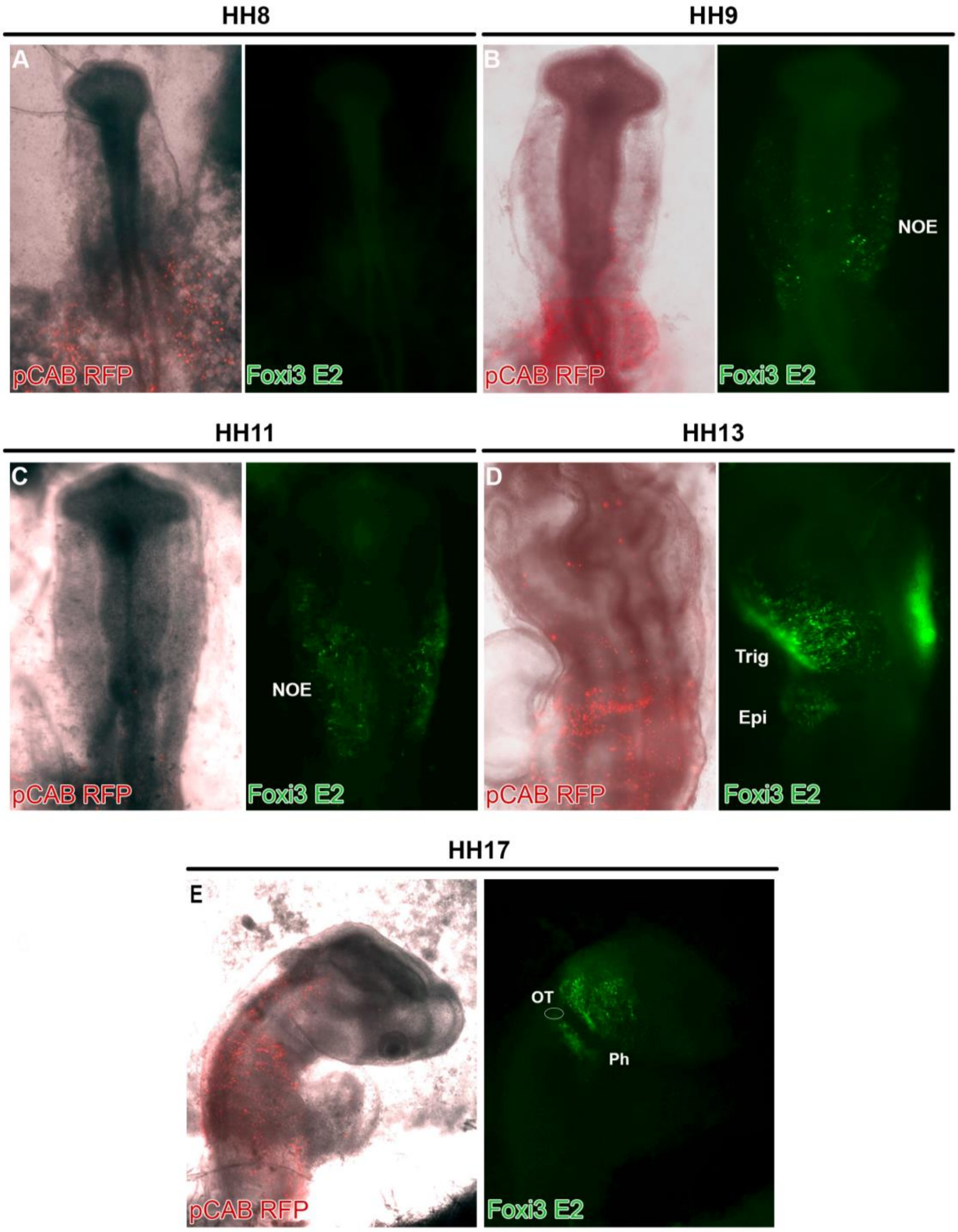


**Supplementary Figure 6.** ***In vivo*** **activity of the Foxi3-E2 enhancer.**
Foxi3-E2 driving eGPF and b-actin driving RFP were coelectroporated into chick embryos at primitive streak stages, and enhancer activity monitored from 0-1ss onwards. Foxi3 E2 GFP+ cells were first identified at HH9 in the Non-Otic Ectoderm (NOE) (B, C) and later found in the trigeminal and epibranchial regions (Trig; Epi) (D). At around HH17 the enhancer remains active in the pharyngeal arches (Ph). The otic vesicle (Ot; circle) has been widely electroporated but no reporter activity is detected (E). Overall the activity of Foxi3 E2 recapitulates the late expression of Foxi3.

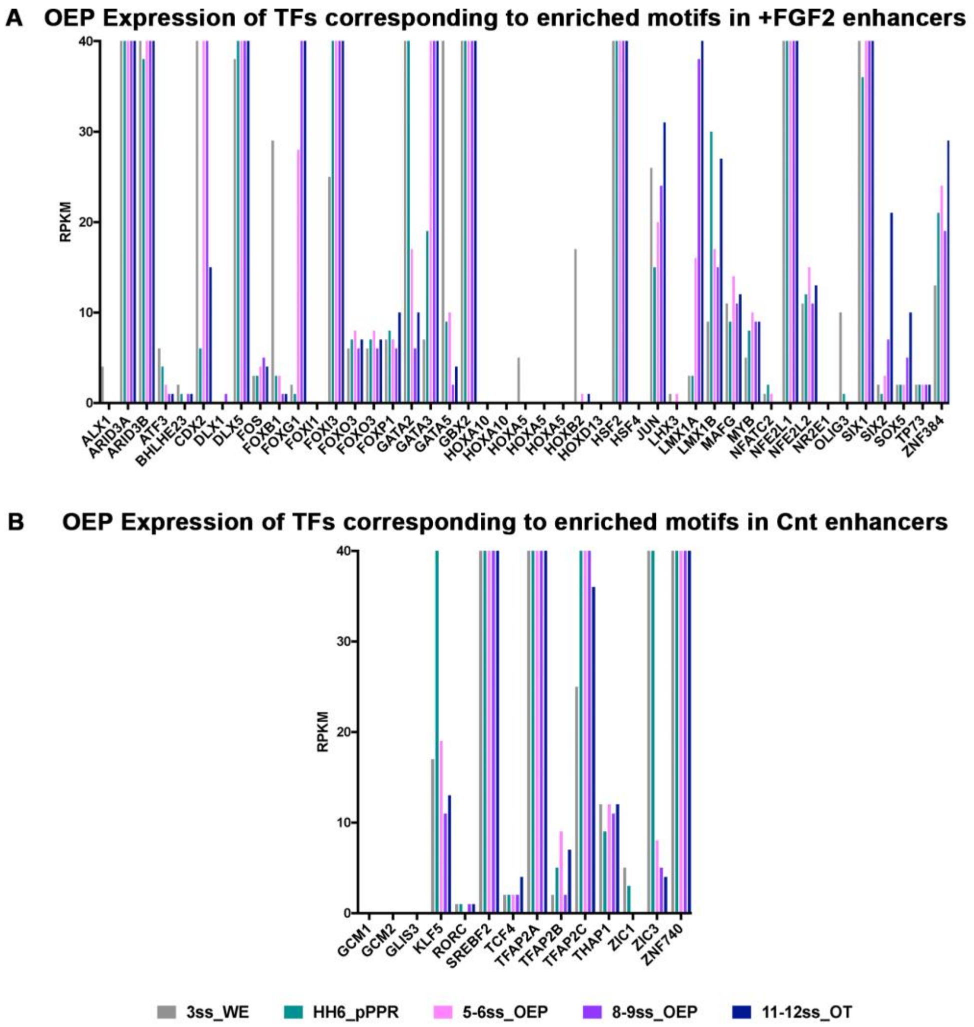


**Supplementary Figure 7. Otic expression of transcription factors corresponding to enriched enhancer motifs.**

Expression level in 3ss whole embryo (grey), 0ss HH6 pPPR (green), 5-6ss OEP (pink), 8-9ss OEP (violet) and 11-12ss otic placode (blue) from Chen and colleagues (Chen et al., 2017) was plotted for transcription factors corresponding to enriched motifs in +FGF2 (A) and control (B) enhancers.

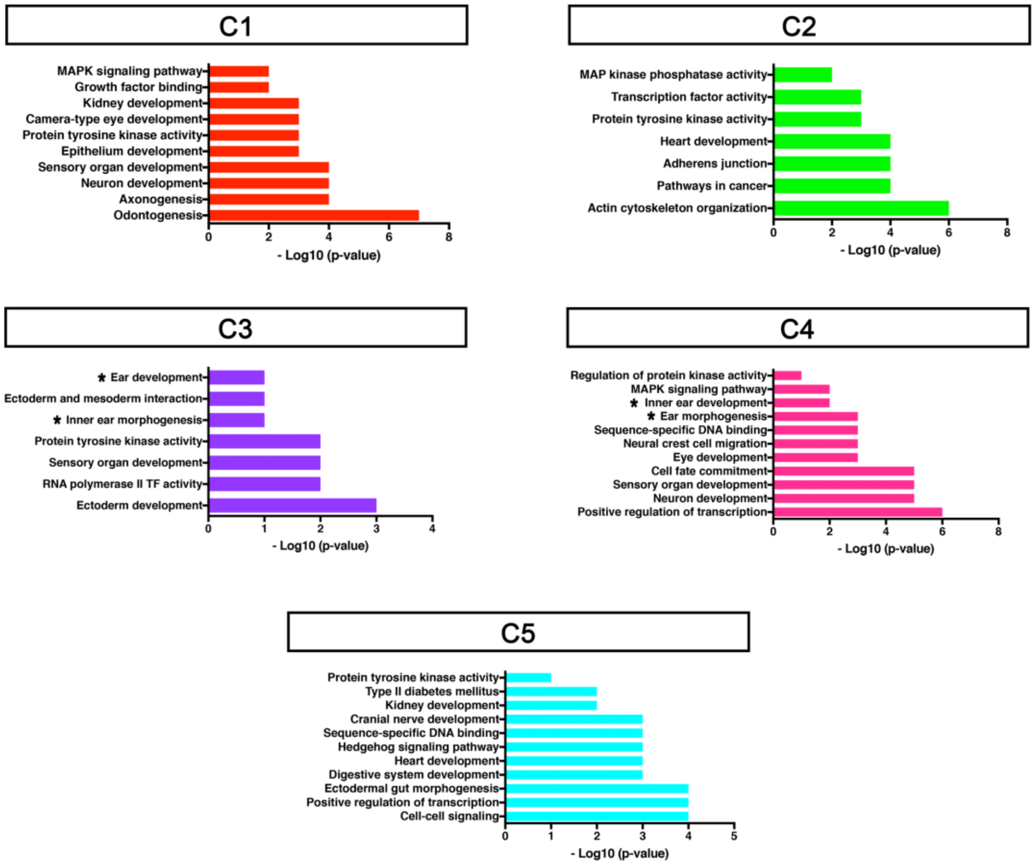


**Supplementary Figure 8. Gene ontology for genes associated with dense H3K27ac peaks flanking Ap1 binding sites.**

To assess each cluster Gene Ontology (GO) and KEGG terms were identified using DAVID. GO terms are coloured according to the colours of each cluster in Figure 3. Cluster 3 and 4 contain putative enhancers for some of the earliest FGF-target genes during OEP induction hence the enriched terms: inner ear morphogenesis (*) and MAPK signalling.

**Supplementary Table 1. Enhancer cloning Primers**

| Enhancer Cloning Primers | | |
|---|---|---|
| Spry1 E1 | F 5'-CTGCCAGCTGTTTCCATTTC-3'<br>R 5'-CTGGGCTGCATGTTGTATTTC-3' | Chr4: 52750797- 52752350<br>1493bp (not active) |
| Spry1 E2 | F 5'-ACGCCTCTCTACCCTCTTT-3'<br>R 5'-GCTGGAAGCTAGAGCCATATC-3' | Chr4: 52768022 - 52768515<br>494bp |
| Foxi3 E1 | F 5'-TCTGACATTTCATCATGGCTTCA-3'<br>R 5'-GGTCATCTGAATGACAACTGTCTC-3' | Chr4: 85595147 - 85595756<br>610bp |
| Foxi3 E2 | F 5'-TTTGGCCCTGTTCAAATGG-3'<br>R 5'-CAGTTTGTTGATACCTTCAGTGT-3' | Chr4: 85611260 – 85611770<br>511bp |
| Hesx1 E | F 5'-GAGACCCTTCAACTTACCAATCT-3'<br>R 5'-GATCCCAGTATCTGAGTGCTTC-3' | Chr12: 8580992 – 8582251<br>1260bp |
| Cxcl14 E | F 5'-AGCCTACCAGTTGTCCTAGA-3'<br>R 5'-CACAGTGTATTGCTTGGCTTT-3' | Chr13: 14642196 -14643846<br>1651bp |

**Supplementary Table 2. ChIP-qPCR Primers**

| P300-Flag ChIP-qPCR Primers | |
|---|---|
| Spry1 E | F 5'-CCTCTATCCCTTTGGTTGTACG-3'<br>R 5'-GATAATGTTTGCTCTGCGGTTC-3' |
| Foxi3 E1 | F 5'-GCAGGGATGGCCTTACATCA-3'<br>R 5'-ACGTGCAGCCATGGAACATA-3' |
| MyoD N | F 5'-AGTCACCTCCACCTAAAATGC-3'<br>R 5'-TGCATGACCGAAGTGTAAGG-3' |

**Supplementary File 1. Cnt_TFBS_Motif_Enrichment**
Summary of RSAT motif enrichment analysis in Cnt enhancer with +FGF2 enhancers as background. Each excel sheet summarises the identified transcription factors for each matrix (Cnt_M1 to Cnt_M30). When no known transcription factor was associated with the enriched motif only k-mer and evalue were reported.

**Supplementary File 2. FGF2_TFBS_Motif_Enrichment**
Summary of RSAT motif enrichment analysis in +FGF2 enhancer with Cnt enhancers as background. Each excel sheet summarises the identified transcription factors for each matrix (Cnt_M1 to Cnt_M30). When no known transcription factor was associated with the enriched motif only k-mer and evalue were reported.

**Supplementary File 3. 3h +FGF2 Nanostring Analysis**

**Supplementary link 4. 6h +FGF2 Nanostring Analysis**
